# The Dynamic Continuum of Nanoscale Peptide Assemblies Facilitates Endocytosis and Endosomal Escape

**DOI:** 10.1101/2021.03.18.435896

**Authors:** Hongjian He, Jiaqi Guo, Jiashu Xu, Jiaqing Wang, Shuang Liu, Bing Xu

**Affiliations:** Department of Chemistry, Brandeis University, 415 South Street, Waltham, MA 02453, USA

**Keywords:** Self-Assembly, Endocytosis, Endosomal Escape, Tissue Nonspecific Alkaline Phosphatase

## Abstract

Considerable number of works have reported alkaline phosphatase (ALP) enabled intracellular targeting by peptide assemblies, but little is known how these substrates of ALP enters cells. Here we show that the nanoscale assemblies of phosphopeptides, as a dynamic continuum, cluster ALP to enable caveolae mediated endocytosis (CME) and eventual endosomal escape. Specifically, fluorescent phosphopeptides, as substrates of tissue nonspecific alkaline phosphatase (TNAP), undergo enzyme catalyzed self-assembly to form nanofibers. As shown by live cell imaging, the nanoparticles of phosphopeptides, being incubated with HEK293 cells overexpressing red fluorescent protein-tagged TNAP (TNAP-RFP), cluster TNAP-RFP in lipid rafts to enable CME, further dephosphorylation of the phosphopeptides form the peptide nanofibers for endosomal escape inside cells. Inhibiting TNAP, cleaving the membrane anchored TNAP, or disrupting lipid rafts abolishes the endocytosis. Moreover, decreasing the formation of peptide nanofibers prevents the endosomal escape. As the first study establishing a dynamic continuum of supramolecular assemblies for cellular uptake, this work not only illustrates an effective design for enzyme responsive supramolecular therapeutics, but also provides mechanism insights for understanding the dynamics of cellular uptakes of proteins or exogenous peptide aggregates at nanoscale.

## INTRODUCTION

Because many drug targets identified by molecular cell biology are inside cells, considerable efforts have focused on engineering molecules for intracellular delivery of various cargo.^1–3^ Besides the use of cationic molecules for enhancing cellular uptake of therapeutics,^4–6^ one of the most explored approaches for intracellular delivery is to engineer molecules to be responsive to chemical or physical stimuli, such as redox,^7–10^ pH,^11–13^ enzymes,^14–23^ or light.^24, 25^ Among enzymatic approach, alkaline phosphatases (ALP) instructed self-assembly of peptides is particularly effective to facilitate the cellular uptake of the peptide assemblies.^14–18, 26–28^ However, the mechanism of this phenomenon is largely unknown. Coincidently, viruses also use responsive motif for cell entry. For example, the life cycle of virus (e.g., SARS-CoV-2)^29^ begins at the attachment to the plasma membrane of host cells^30^ to clustering plasma membrane-bound receptors, followed by viral proteolytic priming (VPP)^31, 32^ for endocytosis and endosomal escape. Prompted by the mechanism of viral cell entry, we hypothesize that the assemblies of phosphopeptides, as the substrate of ALP, act as a dynamic continuum^33^ to cluster the ALP to facilitate endocytosis and to undergo enzymatic morphological transition that enables the endosomal escape (Scheme 1). This hypothesis is also based on several additional rationales: (i) ALP is a GPI-anchor protein, and its clustering enables caveolae mediated endocytosis;^30^ (ii) ALP, locating in lipid rafts, plays a key role in the cellular uptake of aberrant protein aggregates; and (iii) ALP catalyzed dephosphorylation enables morphological transition (e.g., nanoparticles to nanofibers) and generates artificial intracellular filaments.^17^

To prove our hypothesis, we synthesize a phosphopeptide, which, above its critical micelle concentration (CMC), self-assembles to form nanoparticles, which transform into nanofibers upon the dephosphorylation catalyzed by ALP. The phosphopeptide molecules initially aggregate on the plasma membrane to cluster tissue nonspecific alkaline phosphatase (TNAP) in lipid rafts of HEK293 cells that overexpress TNAP, then enable caveolae-mediate endocytosis (CME). The assemblies of the phosphopeptides gradually changes their morphology from nanoparticles to nanofibers depending on the extent of dephosphorylation. Abolishing the interaction between the TNAP and the phosphopeptides significantly decrease the endocytosis. The TNAP, remained in endosome, catalyzes the further dephosphorylation of the phosphopeptides, generating peptide nanofibers to facilitate endosomal escape. As the first study that illustrates the intracellular peptide assemblies resulted from dynamic peptide assemblies rather than a monomeric peptide or static assemblies, this work not only provides insights for understanding the cellular uptakes of proteins or exogenous peptide aggregates, but also offers the guidance for designing enzyme responsive supramolecular assemblies as intracellular targeting therapeutics.

## RESULTS AND DISCUSSION

Scheme 1 shows the structure of the fluorescent phosphopeptide (**NBD-1p**). For studying the cellular uptake of the phosphopeptides, a nitrobenzoxadiazole (NBD), as the fluorophore, conjugates to the side chain of the D-lysine in a naphthyl capped D-phosphotetrapeptide (D-Lys-D-pTyr-D-Phe-D-Phe). Using Fmoc-based solid-phase peptide synthesis, we firstly produced this naphthyl capped D-phosphopeptide, then conjugated it to NBD. Then, the purification by high performance liquid chromatography (HPLC) produces the designed **NBD-1p**. Transmission electron microscopy (TEM) reveals that **NBD-1p** monomers primarily self-assembles into nanoparticles (22.4±7.2 nm in diameter, Figure 1A) at 200 μM (PBS, pH=7.4), which is above its critical micelle concentration (CMC, 159 μM, Figure S1). Upon the addition of ALP for catalytically dephosphorylating **NBD-1p**, the nanoparticles formed by **NBD-1p** transform into short nanofibers (21.9±3.7 nm in diameter and about 300-1000 nm in length) made of **NBD-1** (Figure 1A and Scheme S1). This result confirms that **NBD-1p** undergoes enzyme-instructed self-assembly.^34^

**Scheme 1.**
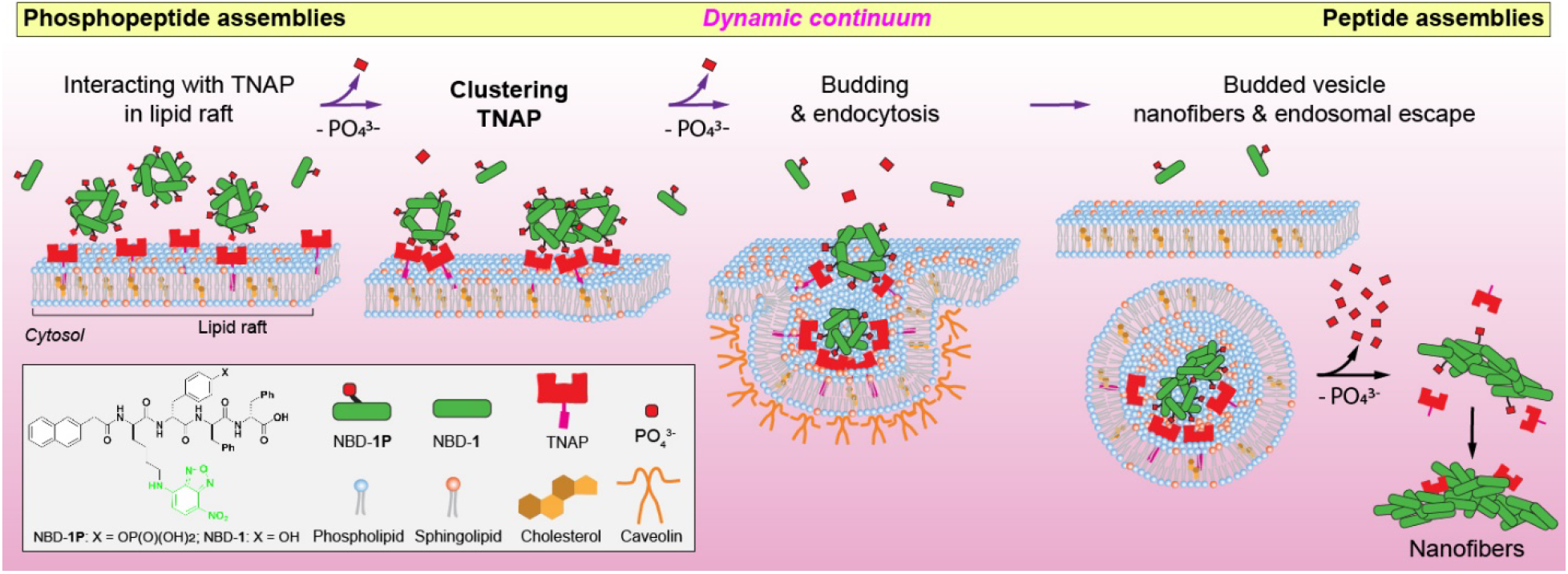
Illustration of a dynamic continuum of peptide assemblies for endocytosis and endosomal escape

**Figure 1.**
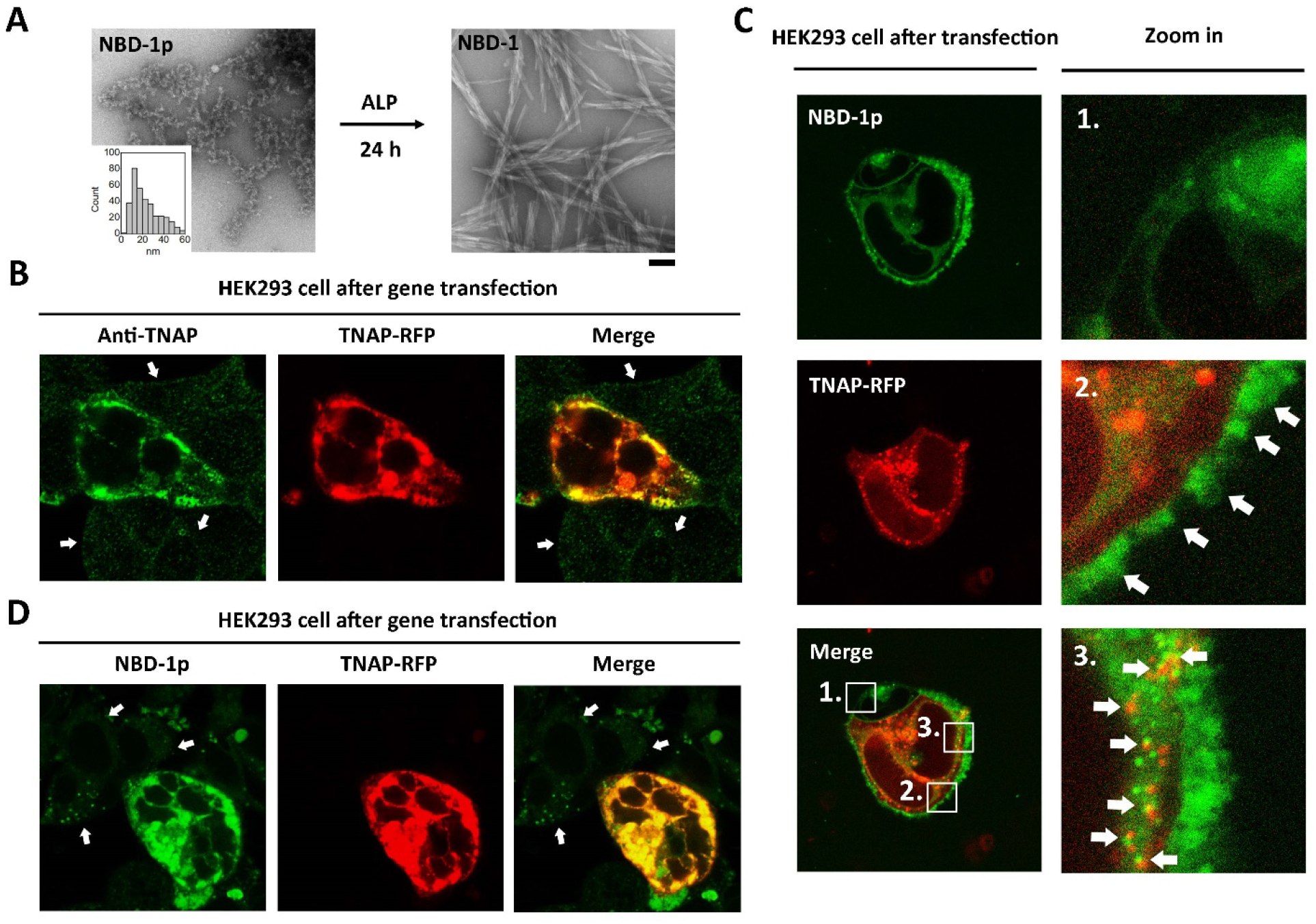
Upregulation of TNAP promotes the endocytosis of phosphopeptide assemblies. (A) TEM images of **NBD-1p** (200 μM) before and after the addition of ALP (0.1 U/mL, 37 °C) for 24 h. The insert is the diameter distribution of the nanoparticles formed by **NBD-1p**. Scale bar = 100 nm. (B) Immunofluorescence staining of TNAP in the transfected HEK293 cell. The non-transfected cells (pointed by arrows) show little TNAP. (C) Confocal fluorescence images of HEK293_TNAP-RFP cells co-incubated with **NBD-1p** (200 μM, 5 min, no washing). To visualize **NBD-1p** at early stage, the laser power here is 5 times stronger than that used in Figure 1D. (D) Confocal fluorescence imaging of TNAP-RFP transfected HEK293 cells treated by **NBD-1p** (200 μM, 12 h, washed). The non-transfected cells (indicated by arrows) show little intracellular NBD fluorescence.

We choose HEK293 cells for testing our hypothesis because of their constitutively low expression of ALP, reliable growth, and propensity for transfection.^35^ Treating HEK293 cells with the polyethyleneimine (PEI)^36^ carrying the DNA plasmid encoding red fluorescent protein-tagged TNAP (TNAP-RFP), we generate a cell population consists of the HEK293 cells that overexpress TNAP-RFP (Figure 1B, the transfected cells are denoted as HEK293_TNAP-RFP) and the HEK293 cells without gene transfection (Figure 1B, indicated by arrows). To observe the fluorescence of **NBD-1p** at the early stage of endocytosis, we used high laser strength for imaging. Co-incubating **NBD-1p** (above CMC) with the HEK293 cells, after the transfection, reveals that, initially (5 min), **NBD-1p** mainly accumulates on the plasma membrane of HEK293_TNAP-RFP cells in a discrete pattern (Figure 1C, region 2, indicated by arrows), although the phosphopeptides barely interact with the plasma membrane of the non-transfected control (Figure 1C, region 1). The cell selective aggregation of **NBD-1p** on the plasma membrane of HEK293_TNAP-RFP cells indicates that the nanoparticles formed by **NBD-1p** associate with the plasma membrane-bound TNAP^37, 38^ for dephosphorylation. The interaction between TNAP and the nanoparticles (made of mainly **NBD-1p** and small amount of **NBD-1**) spatially induces the clustering of TNAP on plasma membrane (Scheme 1). Monomers of **NBD-1p**, being dephosphorylated rapidly, would likely interact with TNAP only briefly. Moreover, we observed not only the formation of more endocytic vesicles in HEK293_TNAP-RFP cells than in the non-transfected control (Figure 1C, region 1 and 3), but also some vesicles containing TNAP-RFP and **NBD-1p** in HEK293_TNAP-RFP cells (Figure 1C, region 3, and Movie S1). These results suggest that the TNAP-participating endocytosis of **NBD-1p** nanoparticles is enhanced in HEK293_TNAP-RFP cells. While some NBD fluorescence exists in some non-transfected HEK293 cells and the endosomal vesicles lacking TNAP in the transfected HEK293 cells, these NBD signals may originate from the cellular uptake of **NBD-1p** via other endocytosis pathways (e. g., macropinocytosis, a nonspecific endocytosis) and the enhanced intracellular NBD fluorescence or autofluorescence under high laser strength. Additionally, **NBD-1p** can still enter the non-transfected HEK293 cells via CME, although inefficiently due to low TNAP expression. The intracellular fluorescence from NBD increases significantly overtime in the HEK293_TNAP-RFP cells, while keeping mostly unchanged in the non-transfected HEK293 cells (Figure 1D, S2 and Movie S2). Trypan blue hardly stains the HEK293_TNAP-RFP cells with **NBD-1p** (Figure S3), confirming an intact cell membrane. Additionally, after the incubation with **NBD-1p**, the TNAP gene transfected HEK293 cells exhibit lower cell viability than that of the wild type (Figure S4). These results confirm that the overexpression of TNAP in HEK293 cell promotes the cellular uptake of the assemblies of **NBD-1p**.

Further study on the endocytosis of **NBD-1p** nanoparticles shows that pretreating the HEK293 cells (after TNAP transfection) with methyl-β-cyclodextrin (M-β-CD), a CME inhibitor via dissipating lipid rafts on plasma membrane,^39, 40^ at the cell compatible concentration (2.5 mM) drastically reduces the cellular uptake efficiency of **NBD-1p** (Figure 2A) compared to the control (Figure 1D). This result indicates that (i) M-β-CD decreases the CME efficiency at that working concentration, and (ii) CME contributes to the cell entry of **NBD-1p** which adheres on the plasma membrane of HEK293_TNAP-RFP cells (Figure 1C, region 2). Since TNAP is enriched in the lipid rafts on plasma membrane,^41^ this result also suggests that the assemblies of **NBD-1p** on plasma membrane (Figure 1C, region 2) associate with the TNAP in lipid rafts, and then cluster the TNAP, which signals the endocytosis of **NBD-1p** through CME^30^ (Scheme 1). The CME of **NBD-1p** nanoparticles triggered by TNAP aggregation agrees with the formation of endocytic vesicles containing TNAP-RFP and **NBD-1p** (Figure 1C, region 3). Using 2,5-dimethoxy-N-(quinolin-3-yl)-benzenesulfonamide (DQB), a noncompetitive inhibitor of TNAP with an allosteric inhibition mechanism,^42^ to inhibit the enzymatic reaction (i.e., TNAP catalyzing the dephosphorylation of **NBD-1p**) or the removal of plasma membrane-bound TNAP via phospholipase C (PLC) which cleaves the GPI anchors^43^ that link TNAP to plasma membrane also substantially decrease the cellular uptake of **NBD-1p** into HEK293_TNAP-RFP cells (Figure 2B and 2C). These results further support that the interaction between TNAP and the nanoparticles of **NBD-1p** is important for signaling CME. Below the CMC, the individual **NBD-1p** molecules neither aggregate on the plasma membrane of HEK293_TNAP-RFP cells (Figure S5 and Movie S3) nor enter the cells efficiently (Figure 2D), although the monomeric phosphopeptides may still associate with the TNAP on cell surface as the substrate of TNAP (Figure 2E). These results indicate that the clustering of TNAP in lipid rafts by the nanoparticles formed by the self-assembly of **NBD-1p** is essential for signaling the CME of the assemblies of **NBD-1p** (Figure 2E).

**Figure 2.**
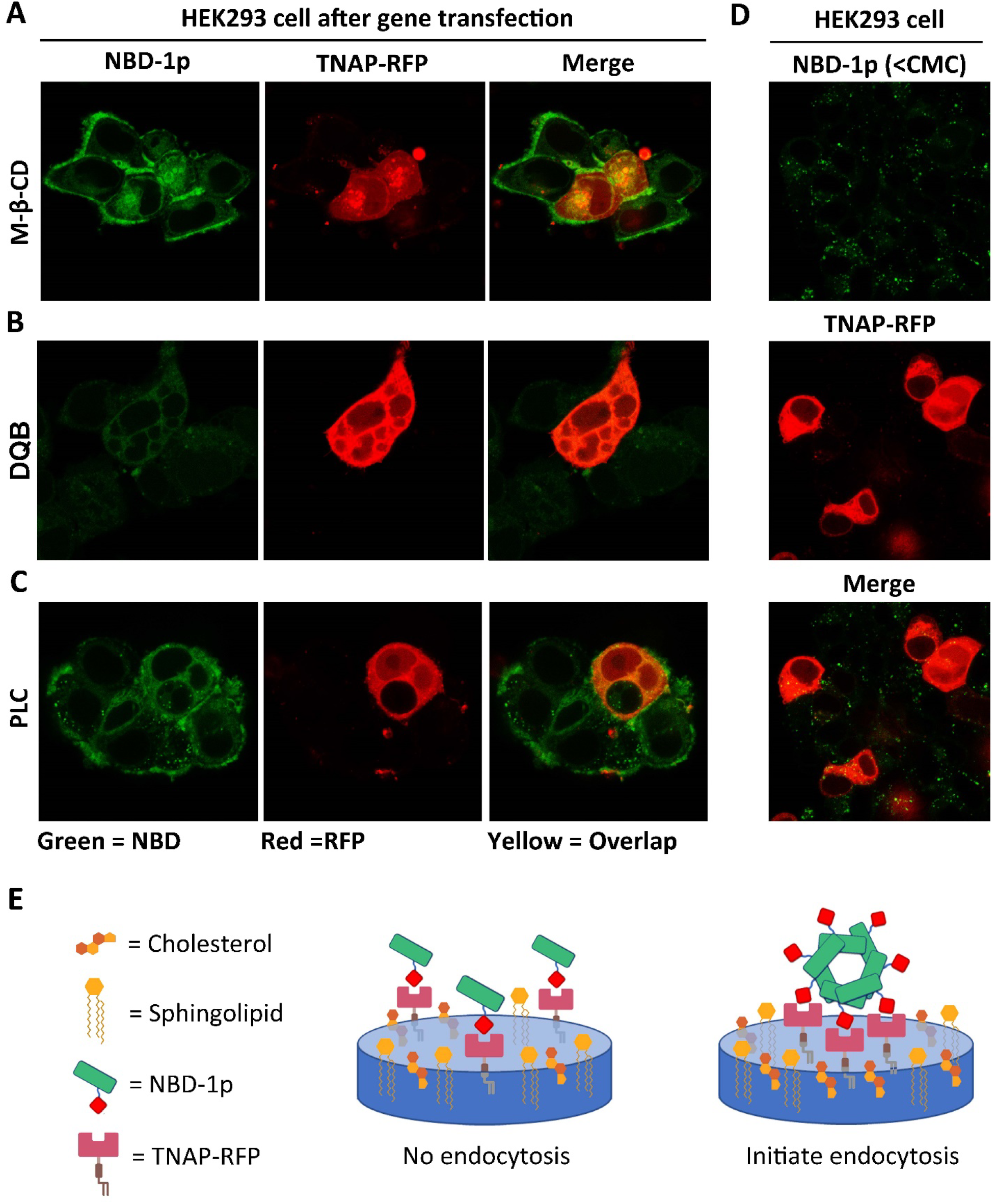
**NBD-1p** nanoparticles associate with the TNAP in lipid rafts followed by CME. (A-C) Confocal fluorescence images of HEK293_TNAP-RFP cells incubated with **NBD-1p** (200 μM, 12 h) after the pretreatment of (A) caveolin-dependent endocytosis inhibitors (M-β-CD), (B) TNAP inhibitor (DQB), and (C) phospholipase C which removes the GPI-anchored TNAP on plasma membrane. (D) Confocal fluorescence images of HEK293_TNAP-RFP cells incubated with **NBD-1p** (50 μM, 12 h) below CMC. (E) Illustration of that the assemblies of phosphopeptide rather than individual molecules initiate CME.

To further validate that the interactions between TNAP and the (D)-pTyr in the nanoparticles of **NBD-1p** initiates the CME, we incubated the HEK293_TNAP-RFP cells with **NBD-1** (i.e., the dephosphorylated **NBD-1p**, Scheme S1). Clearly, the HEK293_TNAP-RFP cells treated by **NBD-1** exhibit significantly less intracellular fluorescence intensity than those treated by **NBD-1p** (Figure 3A and 3B). This result suggests that the interaction between TNAP and the D-pTyr plays a role in the CME of the **NBD-1p** assemblies. Replacing the D-pTry by D-pSer, we generated a derivative of **NBD-1p**, named **NBD-(D)Sp**. The HEK293_TNAP-RFP cells incubated with **NBD-(D)Sp** show drastically reduced cellular uptake of the phosphopeptide compared to the incubation with **NBD-1p** (Figure 3C). Dephosphorylation assay reveals that ALP dephosphorylates **NBD-1p** much faster than **NBD-(D)Sp** at the same concentration (Figure 3D), indicating that ALP exhibits higher affinity for **NBD-1p** than **NBD-(D)Sp**. The results above suggest that the interaction between ALP and its substrate is necessary for the CME of the phosphopeptides. Moreover, a sequence-scrambled phosphopeptide derivative of **NBD-1p**, denoted as **NBD-2p** (with the sequence of D-Phe-D-Phe-D-Lys(◻-NBD)-D-pTyr, Scheme S1), also enters the HEK293_TNAP-RFP cells more efficiently than the non-transfected HEK293 cells (Figure S6). This result indicates that the cellular uptake of the phosphopeptide assemblies containing D-pTyr depends more on the interactions between TNAP and the enzyme triggers than on the binding between receptors and specific peptide sequences.^44–46^

**Figure 3.**
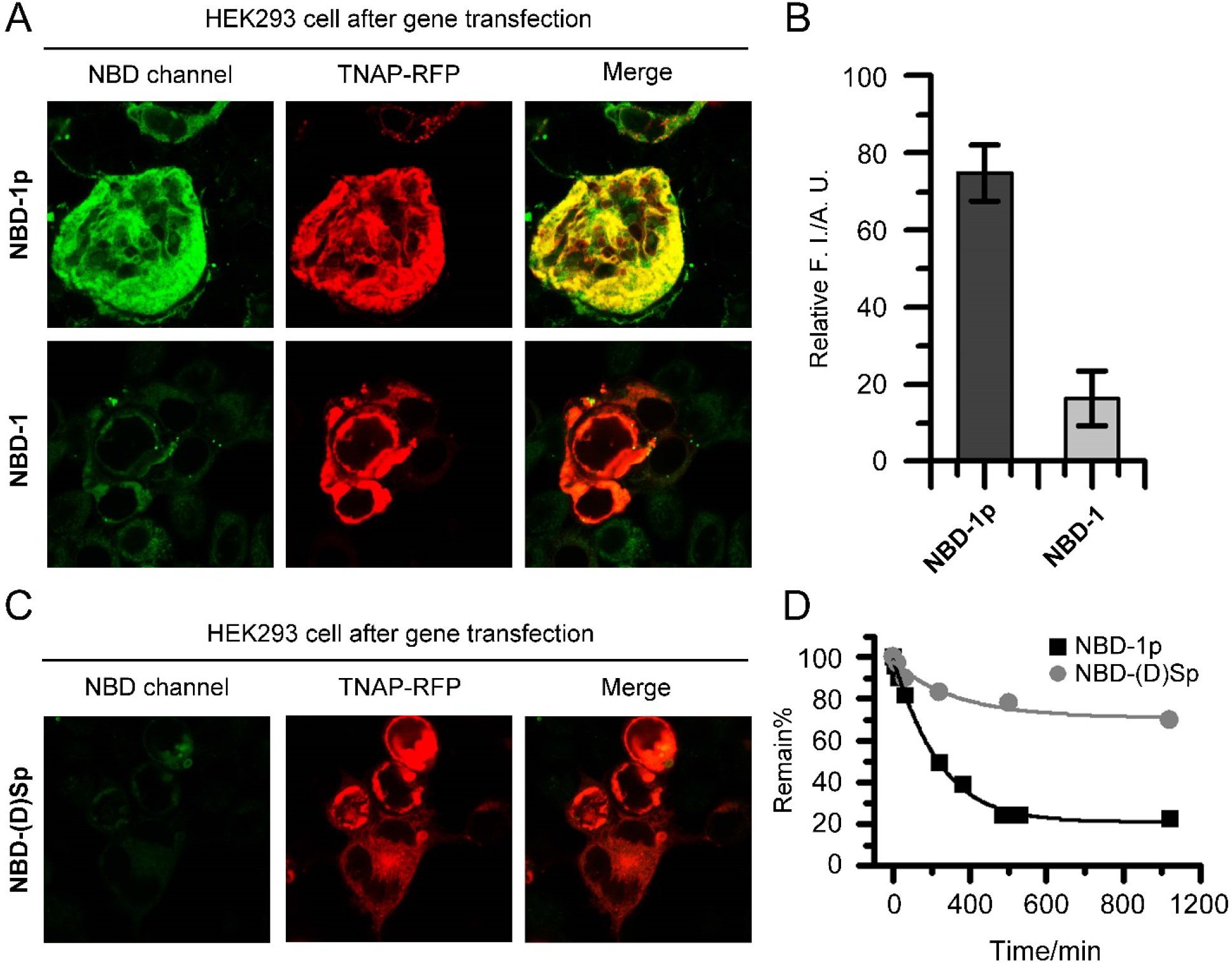
Phosphotyrosine is critical for rapid cellular uptake. (A) Confocal fluorescence images of HEK293_TNAP-RFP cells incubated with **NBD-1p** or **NBD-1** (200 μM, 12 h, washed), respectively. (B) ImageJ analysis of intracellular fluorescence intensity of HEK293_TNAP-RFP cells in (A). Data is presented as mean ± standard deviation. (C) Confocal fluorescence images of HEK293_TNAP-RFP cells incubated with **NBD-(D)Sp** (400 μM, 12 h, washed). (D) Dephosphorylation rate of **NBD-1p** and **NBD-(D)Sp** by ALP (0.1 U/mL, 37 °C).

The dispersive fluorescence of **NBD-1p** in HEK293_TNAP-RFP cells (Figure 1D and 3A) not only confirms the endocytosis of **NBD-1p**, but also the endosomal escape of the peptide assemblies. Considering the cell entry mechanism of SARS-CoV-2, in which the proteolytic cleavage of S proteins by endosomal proteases (e.g., cathepsin B and cathepsin L) promotes the release of viral genome from endosome to cytosol,^47^ we reckon that, after the CME of the assemblies of **NBD-1p**, further dephosphorylation catalyzed by the TNAP in endosomal compartments induces a phase transition on the peptide assemblies, which facilitates the endosomal escape of NBD-1 into cytosol. Liquid chromatography-mass spectrometry (LC-MS) analysis of the cell lysate of HEK293_TNAP-RFP cells that are preincubated with **NBD-1p** confirms the dephosphorylation of the peptide inside cells (Figure S7). Additionally, TEM images of liposomes encapsulating **NBD-1p** and ALP exhibit the generation of nanofibers that rupture the liposomes (Figure 4A). These results imply that the TNAP in endocytic vesicles convert the **NBD-1p** nanoparticles to nanofibers (**NBD-1**), which break the membrane of endocytic vesicles for escaping into cytosol. More TEM images reveal that the **NBD-1p** assemblies mixed with ALP initially remain as nanoparticles until certain dephosphorylation ratio that triggers the particle-to-fiber phase transition (Figure S8 and 3D). This result implies that although dephosphorylation begins once TNAP binds to **NBD-1p** assemblies on plasma membrane, the assemblies likely maintain as nanoparticles (which consist of mainly **NBD-1p** and small amount **NBD-1**) during internalization until being further dephosphorylated in endosomal vesicles where nanofibers form for endosome rupturing. This observation also agrees with the reports that filamentous nanostructures have much reduced/limited cellular uptake ability relative to their spherical counterparts.^48, 49^

**Figure 4.**
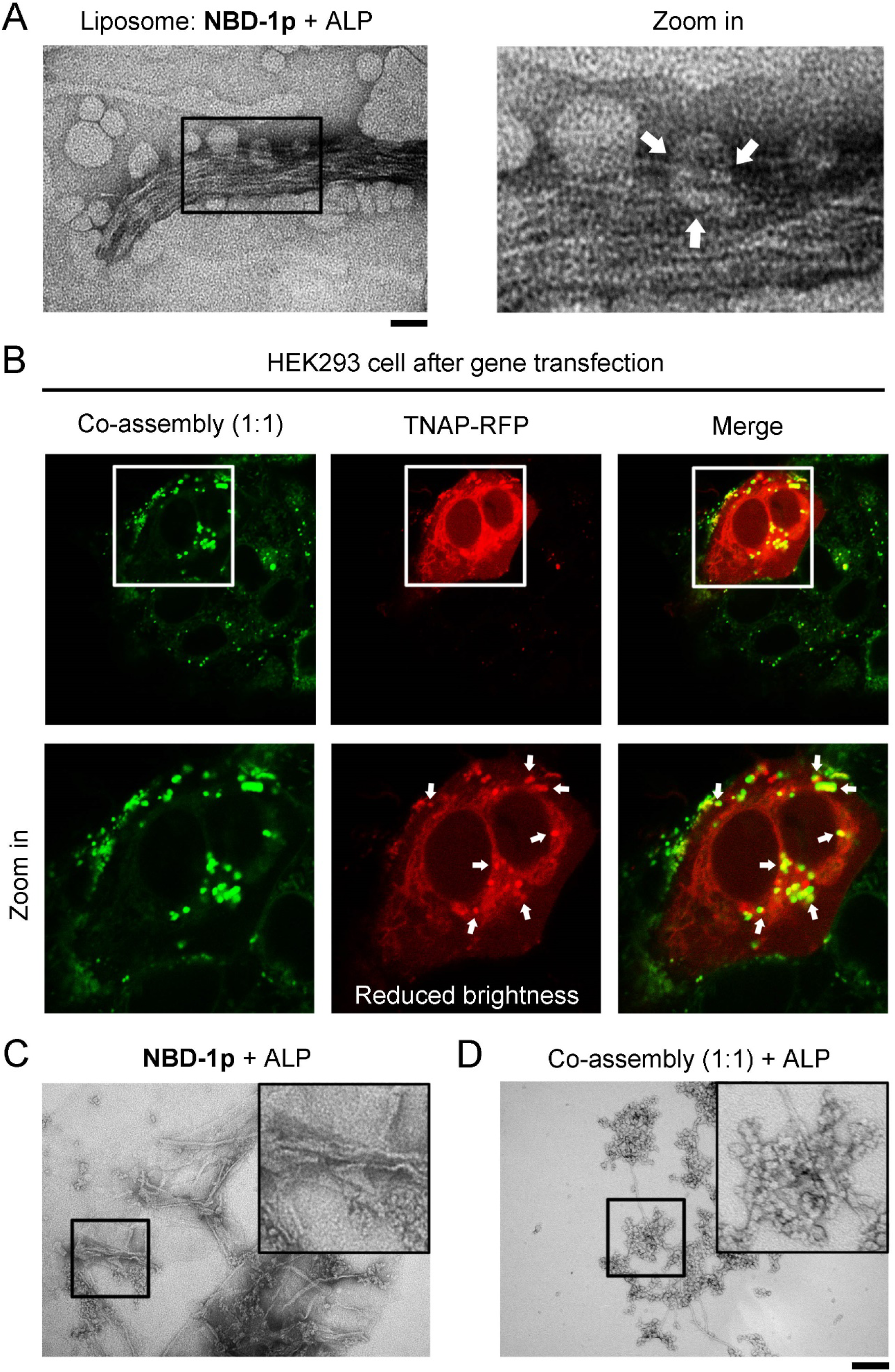
The co-assembly of **NBD-1p** and **NBD-(D)Sp** shows less efficient endosomal escape. (A) TEM image of liposomes carrying **NBD-1p** and ALP (1 U/mL, 37 ◻, 6 h). Broken liposomes are indicated by arrows. Scale bar = 100 nm. (B) Confocal fluorescence images of HEK293_TNAP-RFP cells incubated with the mixture of **NBD-1p** (100 μM) and **NBD-(D)Sp** (100 μM) for 12 h. The peptides trapped in endosome are indicated by arrows. (C) TEM images of **NBD-1p** (200 μM) and (D) the co-assemblies of **NBD-1p** (100 μM) and **NBD-(D)Sp** (100 μM) after the addition of ALP (0.1 U/mL, 37 ◻, 3 h) for partial dephosphorylation. ALP yields more nanofibers in (C) than in (D). Scale bar = 100 nm.

To further prove the above assumption of endosomal escape, we incubate HEK293 cells, after TNAP-RFP gene transfection, with the mixture of **NBD-1p** and **NBD-(D)Sp** (total peptide concentration is 200 μM). The HEK293_TNAP-RFP cells incubated with the mixture of **NBD-1p** and **NBD-(D)Sp** at 1:1 or 1:3 ratio mainly generates fluorescent puncta colocalizing with TNAP-RFP inside the cells (Figure 4B and S9A), although the cells incubated with **NBD-1p** (100 or 50 μM) or **NBD-(D)Sp** (100 or 150 μM)), respectively, below CMC exhibit little fluorescence (Figure S10). This result indicates that **NBD-1p** and **NBD-(D)Sp** form co-assemblies in solution, and the co-assemblies of **NBD-1p** and **NBD-(D)Sp** (1:1 or 1:3) primarily retain in the endocytic vesicles containing TNAP-RFP after the endocytosis. Our results also indicate that phosphopeptide assemblies exhibits emergent properties, which are absent in monomeric phosphopeptides. The nanoparticles formed by **NBD-1p** transform into nanofibers for endosomal rupturing after partial dephosphorylation by ALP (Figure 4C and 3D). Thus, the remained phosphopeptides, after releasing into cytosol, get further dephosphorylation by intracellular TNAP. This process produces networks of peptidic nanofibers that encapsulate intracellular TNAP (Figure 1D and 3A, TNAP colocalizes with NBD). However, the co-assemblies of **NBD-1p** and **NBD-(D)Sp** (1:1 or 1:3) mixed with ALP mostly remain as nanoparticles (Figure 4D and S9B) because **NBD-(D)Sp** is a poor substrate of ALP (Figure 3D). When the mixture has the ratio of **NBD-1p** and **NBD-(D)Sp** in 3 to 1, the co-assemblies behave similarly to the assemblies of **NBD-1p** (Figure S9). Moreover, a phosphopeptide derivative (**NBD-3p**) carrying D-pTry and pSer (Scheme S1) also ends up in the endosomal compartments of HEK293_TNAP-RFP cells after incubation (Figure S11). Like the co-assemblies of **NBD-1p** and **NBD-(D)Sp** with 1:1 or 1:3 ratio, **NBD-3P**, being treated by ALP, inefficiently undergoes the nanoparticle-to-nanofiber transition (Figure S12). These results suggest that the particle-to-fiber transition of the phosphopeptide assemblies catalyzed by the ALP in endosome is critical for the endosomal escape of the peptide assemblies, likely via rupturing the endosome by the formation of nanofibers.

## CONCLUSIONS

In conclusion, this work demonstrates the continuous morphological transformation of peptide assemblies, catalyzed by ALP, is responsible for the endocytosis, mainly through caveolae-mediated endocytic pathway, and endosomal escape of the peptide assemblies. Although other endocytosis pathways, such as macropinocytosis, may contribute to the cellular uptake of phosphopeptide assemblies, they are less likely act as the main path for the phosphopeptide assemblies (Figure S13). Here, TNAP catalytically controls a dynamic morphological transition, thus enables endocytosis and endosomal escape of peptide assemblies, in an analogy to the cell entry of virus. Because **NBD-1p** mainly exists as nanoparticles at the concentration above CMC, the effects of monomeric **NBD-1p** likely are insignificant. We suggest this type of cell uptake as “Enzyme Primed Endocytosis (EPE)”. While the contribution of VPP in the cell entry of virus has been extensively studied, little research elucidates how EPE controls the cellular uptake of other molecular assemblies. In analogy to the ligand-receptor mediated endocytosis that require tight binding, the clustering of enzymes in lipid rafts and the enzymatic reaction of supramolecular assemblies require rapid enzyme kinetics because **NBD-(D)Sp** hardly enter the cells. This study also provides a mechanistic understanding of the role of peptide assemblies for context-dependent signaling.^50, 51^ Although this work focuses on ALP and peptides, the mechanism revealed likely is applicable for the endocytosis of the substrates of esterase^52^ or proteases,^53^ including non-peptide substrates of enzymes.

## Supporting information

ESI

## ASSOCIATED CONTENT

### Supporting Information

The Supporting Information is available free of charge at: https://pubs.acs.org/doi/xxx.

Experimental Procedures; (Scheme S1) Molecular structure; (Figure S1) Critical micelle concentration; (Figure S2) Time-dependent endocytosis; (Figure S3) Trypan blue staining of cells; (Figure S4) MTT cell viability assay; (Figure S5) Confocal fluorescence images of the HEK293 cells after TNAP-RFP gene transfection incubated with phosphopeptide below CMC; (Figure S6) Confocal fluorescence imaging of HEK293 cells after the transfection of TNAP-RFP gene incubated with sequence-scrambled phosphopeptide; (Figure S7) LC-MS analysis of cell lysate of HEK293 cells; (Figure S8) TEM images of phosphopeptide incubated with ALP for different time; (Figure S9) HEK293 cells, after TNAP-RFP gene transfection, incubated with the co-assemblies of **NBD-1p** and **NBD-(D)Sp**; (Figure S10) HEK293 cells, after TNAP-RFP gene transfection, incubated with **NBD-1p** or **NBD-(D)Sp** below CMC, respectively; (Figure S11) Confocal fluorescence imaging of HEK293 cells, after the transfection of TNAP-RFP gene, incubated with **NBD-3p**; (Figure S12) TEM images of **NBD-3p** incubated with ALP; (Figure S13) Confocal fluorescence images of HEK293_TNAP-RFP cells incubated with **NBD-1p** in the presence of macropinocytosis inhibitor; and (Figure S14) Mass spectrum.

### Author Contributions

B.X. and H. H. designed the study. H. H. performed the experiment and generated the data. J. G. and J. X. assisted in the synthesis of peptides. J. W. and S. L. assisted in cell culture. B. X. and H. H. wrote the manuscript.

### Funding Sources

NIH (R01CA142746) and NSF (DMR-2011846).

### Notes

The authors declare no competing financial interest.

## ACKNOWLEDGMENT

The authors thank NIH (R01CA142746) and NSF (DMR-2011846) for their support.

