## Supplementary material for "The Dynamic Continuum of Nanoscale Peptide Assemblies Facilitates Endocytosis and Endosomal Escape": ESI

Experimental Procedures

**Materials**

All amino acid derivatives involved in the synthesis were purchased from GL Biochem (Shanghai) Ltd. N, N-diisopropylethylamine (DIEA) and O-benzotriazole-N,N,N',N'-tetramethyluronium-hexafluorophosphate (HBTU) were purchased from Fisher Scientific. The synthesis of all peptide fragments was based on solid-phase peptide synthesis (SPPS). 4-Chloro-7-nitrobenzofurazan (NBD-Cl) was purchased from Tokyo Chemical Industry (TCI). The peptides were made *via* the combination of SPPS and solution phase synthesis. All crude compounds were purified by HPLC with the yield of 70-80%. All reagents and solvents were used as received without further purification unless otherwise stated. All cell lines were purchased from ATCC. All media for cell culture were purchased from ATCC, fetal bovine serum (FBS) and penicillin/streptomycin from Gibco by Life technologies, and 3-(4,5-Dimethylthiazol-2-yl)-2,5-diphenyltetrazolium bromide (MTT) from ACROS Organics. All antibodies were purchased from Abcam. Alkaline phosphatase was purchase from Thermo Fisher Scientific. TNAP Inhibitor (DQB) was purchased from EMD Millipore.

**Instruments**

All peptides were purified by Water Agilent 1100 HPLC system, equipped with an XTerra C18 RP column. LC-MS was operated on a Waters Acquity Ultra Performance LC with Waters MICRO-MASS detector. Transmission electron microscope (TEM) images were taken on Morgagni 268 transmission electron microscope. Fluorescent analysis was performed on Shimadzu RF-5301-PC fluorescence spectrophotometer. Fluorescence images were taken by ZEISS LSM 880 confocal laser scanning microscope. Circular dichroism (CD) spectra were obtained by Jasco J-810 spectropolarimeter.

Experimental Procedures

**Synthesis of NBD-labeled Peptides**. The synthesis of all peptides was done by Fmoc-based solid phase synthesis^[1]^. 1 mmol of NBD-Cl was dissolved in methanol. The peptides (1 mmol) were dissolved in water. 1.5 mmol Na_2_CO_3_ was added into the solution of peptides. A clear aqueous solution of peptide was observed after the addition of Na_2_CO_3_. NBD-Cl in methanol was added into the solution of peptide by drops. Thin layer chromatography (TLC) was used to trace the reactions. HPLC was used to purify the final products.

**Gene Transfection**. The gene transfection in HEK293 cells was done according to a previously reported method.^[2]^ Briefly, Alkaline Phosphatase cDNA ORF Clone, Human, C-OFPSpark tag gene (Sino Biological Inc., Cat: HG10440-ACR) was incubated with polyethylenimine (PEI, Mw = 25, 000) for 20 min. Then, the mixture was added to HEK293 cells incubated with serum-free culture medium for another overnight incubation. After the incubation, cells were detached from petri dish by trypsin, and then 1/3 of the cells were seeded backed to the petri dish. The cells were incubated in cell culture hood until further analysis.

**Confocal fluorescence imaging**. Cells were seeded in confocal dishes 24 hours prior to experiment. After being treated by conditions of interest, the cells were washed by PBS for 3 times followed by the imaging using ZEISS LSM 880 confocal laser scanning microscope. The experiment conditions are listed as follow unless specifically mentioned: NBD channel, ex 488 nm, em 500-600 nm, laser power 1%; RFP channel, ex 561 nm, em 570-640 nm, laser power 5%. For imaging the **NBD-1p** at the early stage of endocytosis (Figure 1C and Movie S1), the power of 488 nm laser is 5%. For Movie S2 and S3, the 488 nm laser power is 0.5% and the 561 nm laser power is 3% for reducing the photobleaching and heating effect.

**Determination of Dephosphorylation in Cells**. NBD-1p (200 μM) was incubated with HEK293 cells after the transfection of TNAP-RFP gene. After the incubation, cells were washed by PBS for 3X. Lysis buffer was added to the cells and the cell lysate was collected. Then, the cell lysate was analyzed by LC-MS.

**Liposome preparation**. The liposome kit was purchased from Sigma (Cat: L4395-1VL). Liposome was prepared according to manufacturer’s instruction. The mixture of NBD-1p and ALP (1 U/mL, in PBS) was added to the vial containing liposome component followed by vortex for 30 min. After that, the whole mixture was centrifuged, and the supernatant containing the unencapsulated NBD-1p and ALP was removed. The precipitate was resuspended in PBS and stored in 4 ℃ before further analysis.

**TEM Images**. TEM images were taken on Morgagni 268 transmission electron microscope using negative staining. Following an established protocol^[3]^, samples (dissolved in PBS) were dropped on copper grids and dried. Uranium acetate was used to stain the samples. Images were taken by lab members who were properly trained.

**Cell Culture**. HeLa cells were cultured in Minimum Essential Medium (MEM) supplemented with 10% (vol/vol) FBS, 0.5% (vol/vol) penicillin (10, 000 unit), and 0.5% (vol/vol) streptomycin (10, 000 unit). HS-5 cells were cultured in Dulbecco’s Modified Eagle Media (DMEM) supplemented with 10% (vol/vol) FBS, 0.5% (vol/vol) penicillin (10, 000 unit), and 0.5% (vol/vol) streptomycin (10, 000 unit). Saos-2 cells were cultured in McCoy's 5a medium supplemented with 15% fetal bovine serum, and 0.5% (vol/vol) streptomycin (10, 000 unit). All cells were maintained at 37 °C in a 5% CO_2_ incubator.

**Cell Viability Assay**. The cell viability was determined by MTT assay. Cells were seeded in 96-well plates (1X10^4^ cells/well) and incubated in a cell incubator for overnight for adhesion. After adhesion, culture medium in each well was removed followed by adding different concentration of the compounds of interests (dissolved in culture medium). To examine the cell viability, 10 L MTT solution (5 mg/mL) was added into the wells designed for 1, 2 and 3 days incubation, respectively. The plates were incubated in an incubator for another 4 hours. 100 μL SDS-HCl solution was added to stop the MTT-reduction reaction and to dissolve the formazan. After measuring the absorbance of each well at 595 nm with multimode microplate reader, we calculated the cell viability percentage relative to the untreated cells (solvent control). The MTT assay was performed in triplet, and the average value of the three measurements was taken.

**Semi-Quantification of Intracellular Fluorescence Intensity**. Cells treated by conditions of interest were analyzed by confocal fluorescence microscopy. ImageJ was used to quantify the fluorescence intensity in targeted cells. Finally, the fluorescence intensity was average, and the data was presented as mean ± standard deviation.

**Immunofluorescence Staining**. Cells were plated on confocal dishes (CellVis) and fixed in 4 wt% paraformaldehyde for 15 min and permeabilized with 1% BSA and 0.1% Tween 20. Fixed cells were incubated in primary antibody at 4 ^o^C overnight, washed three times for 5 min each, incubated in secondary antibody for 1 h, then washed three times for 5 min each.

**Quantification of Dephosphorylation Rate**. Phosphopeptides were incubated with ALP in PBS buffer at 37 ^o^C. The mixtures were analyzed by HPLC after the incubation of different time (e.g., 0 min, 10 min, 30 min, 1 h, 2 h, 4 h, 8 h, 24 h, etc.). The remain% was determined by integrating the peak area in HPLC spectra. Remain% = Area [Phosphopeptide]_current_/Area [Phosphopeptide]_0 min_X100%.

Determination of CMC. The CMC of NBD-1p was determined using pyrene as the fluorescent probe^[4]^. Briefly, different concentrations of 1 were prepared in pyrene-saturate solutions. The fluorescence spectra of pyrene solutions with different concentration of NBD-1p were obtained. The intensity ratio of the 1^st^ peak/the 3^rd^ peak (I_1_/I_3_) was determined. Plot I_1_/I_3_ against the concentrations of NBD-1p. The concentration at the turning point is the CMC.

Results and Discussion

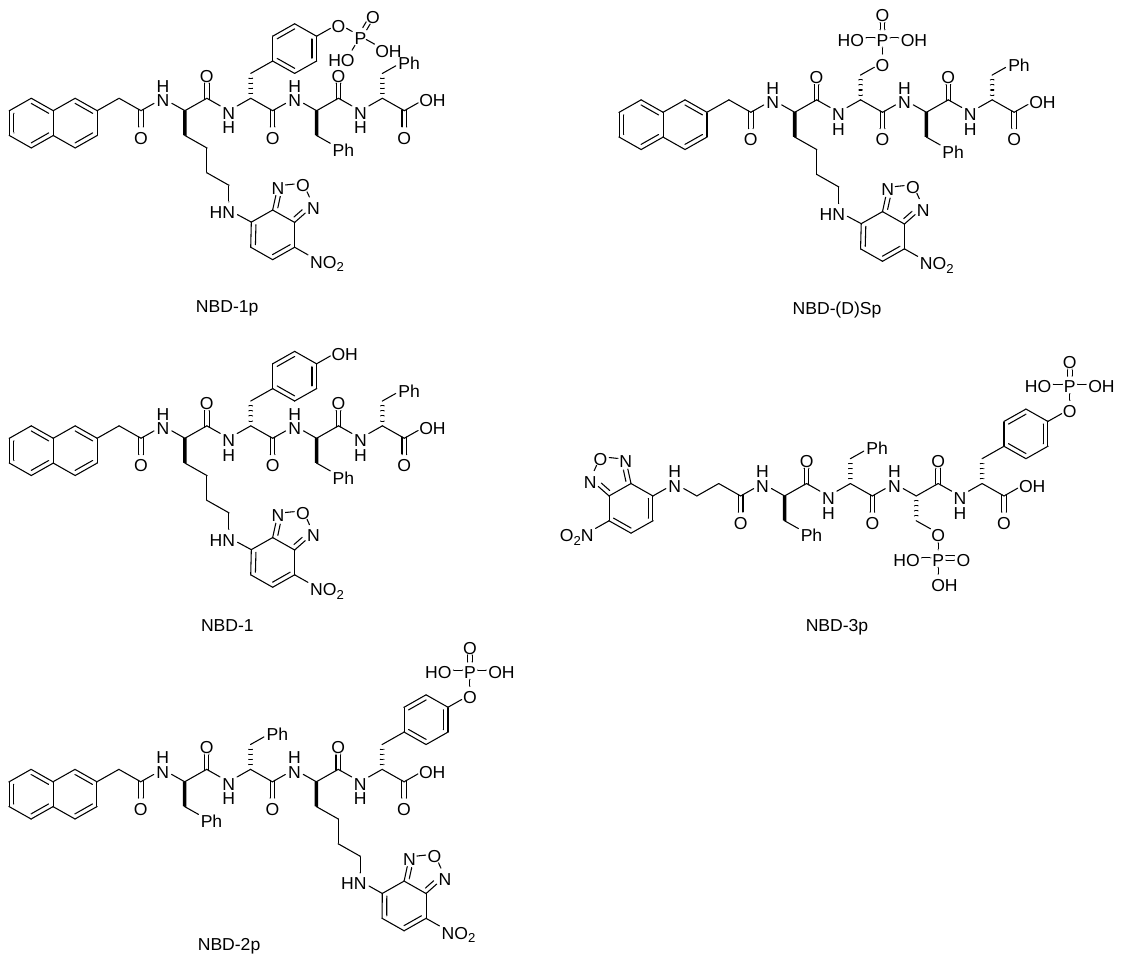

**Scheme S1.** Molecular structure of NBD-1p, NBD-1, NBD-2p, NBD-3p and NBD-(D)Sp.

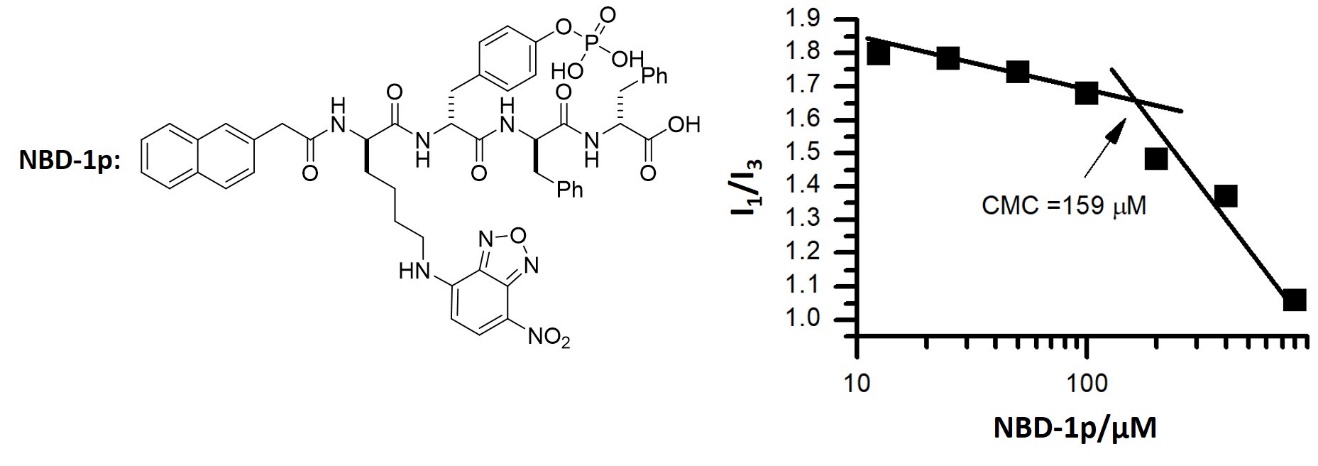

**Figure S1.** Critical micelle concentration of NBD-1p.

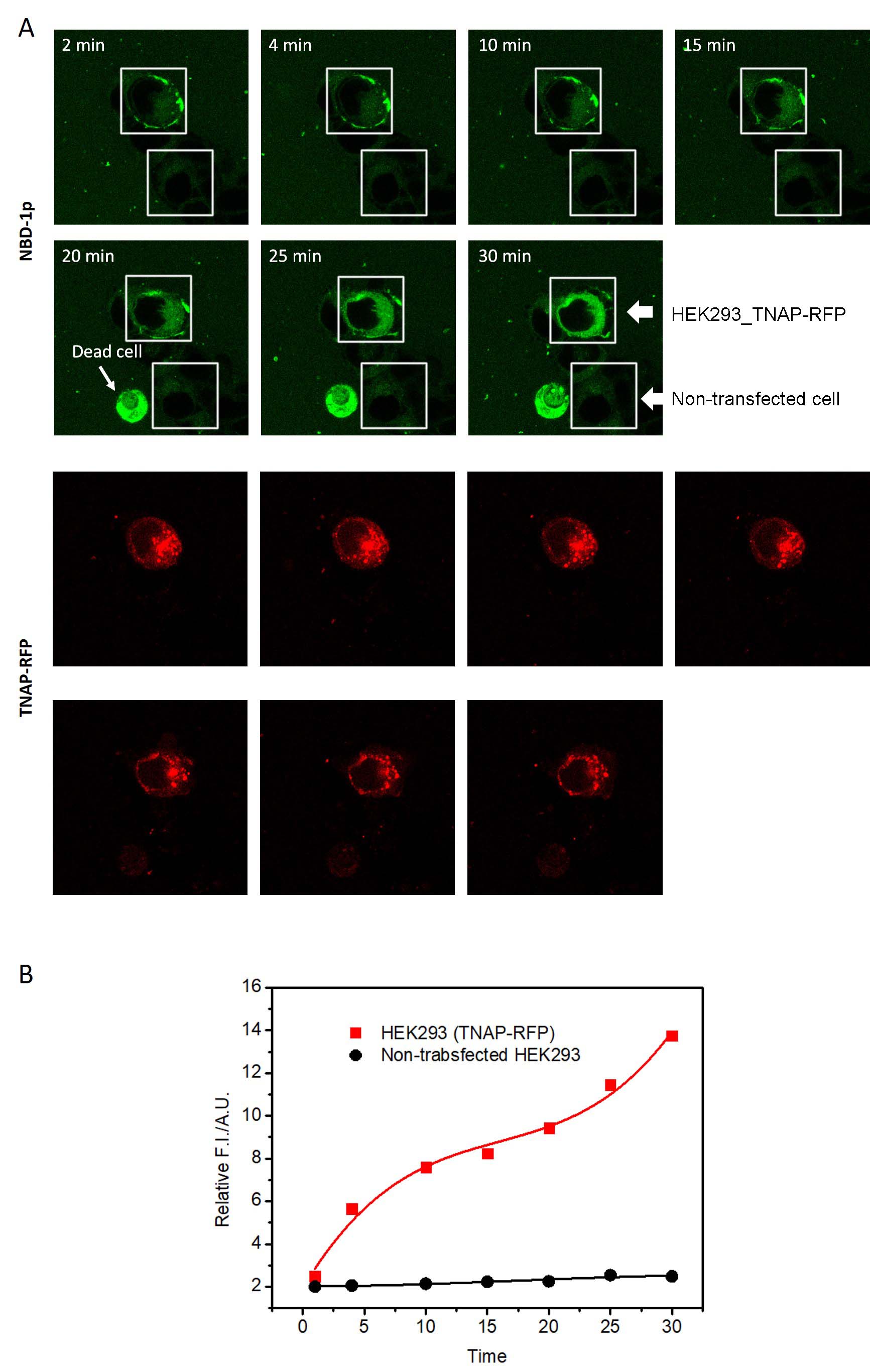

**Figure S2.** Time-dependent endocytosis of NBD-1p. (A) Confocal fluorescence images of the non-transfected HEK293 and HEK293_TNAP-RFP cells incubated with NBD-1p (200 μM) for different time. (B) Semi-quantification of the intracellular intensity of NBD in the non-transfected HEK293 and HEK293_TNAP-RFP cells incubated with NBD-1p (200 μM) for different time.

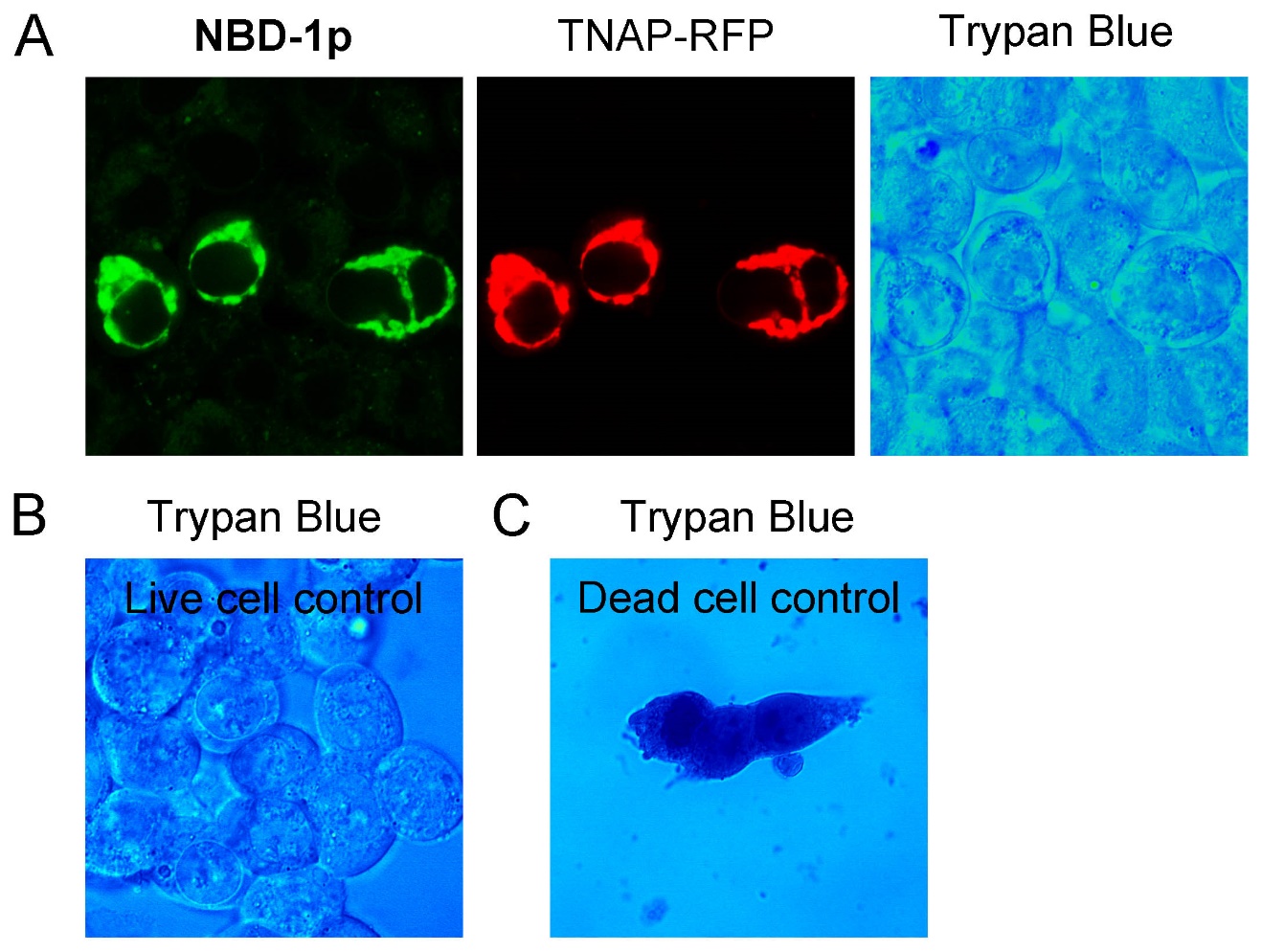

**Figure S3.** Trypan blue staining of HEK293_TANP-RFP cells incubated with (A) NBD-1p (200 μM, 12 h), (B) culture medium only (live cell control), and (C) cisplatin (100 μM, 24 h), which serve as dead cell control.

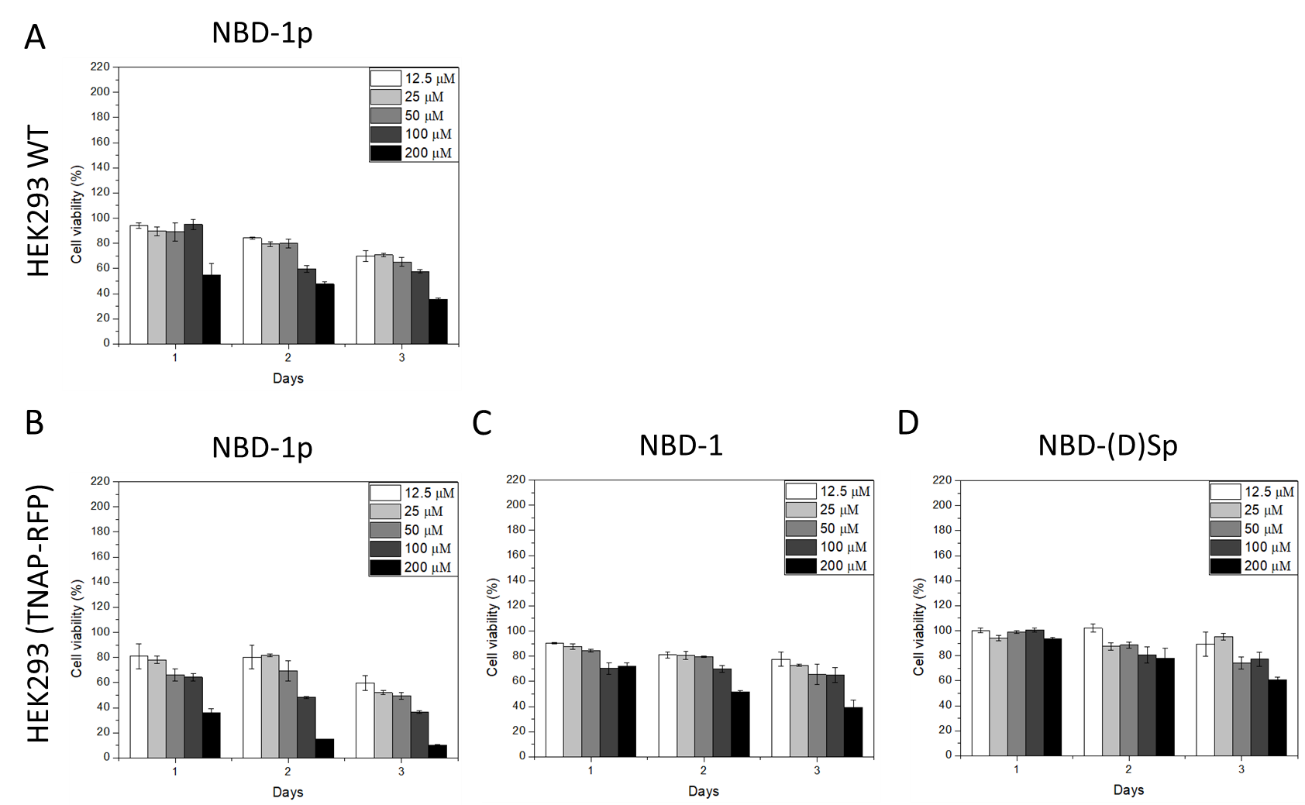

**Figure S4.** MTT cell viability assay of (A) HEK293 (WT) and (B) HEK293 cells after gene transfection in the presence of different concentrations of NBD-1p. (C) Cell viability of HEK293 cells, after gene transfection, incubated with NBD-1 at different concentration. (D) Cell viability of HEK293 cells, after gene transfection, incubated with NBD-(D)Sp at different concentration.

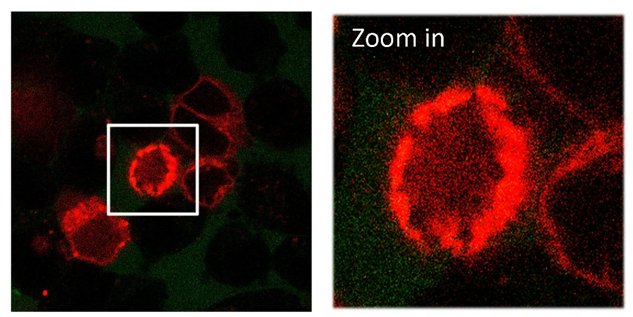

**Figure S5.** Confocal fluorescence images of the HEK293 cells, after TNAP-RFP gene transfection, co-incubated with NBD-1p (50 μM, 5 min, without washing) below CMC. No assembly of NBD-1p is observed on the plasma membrane of HEK293_TNAP-RFP cells at this concentration. Cells were not washed by PBS before imaging.

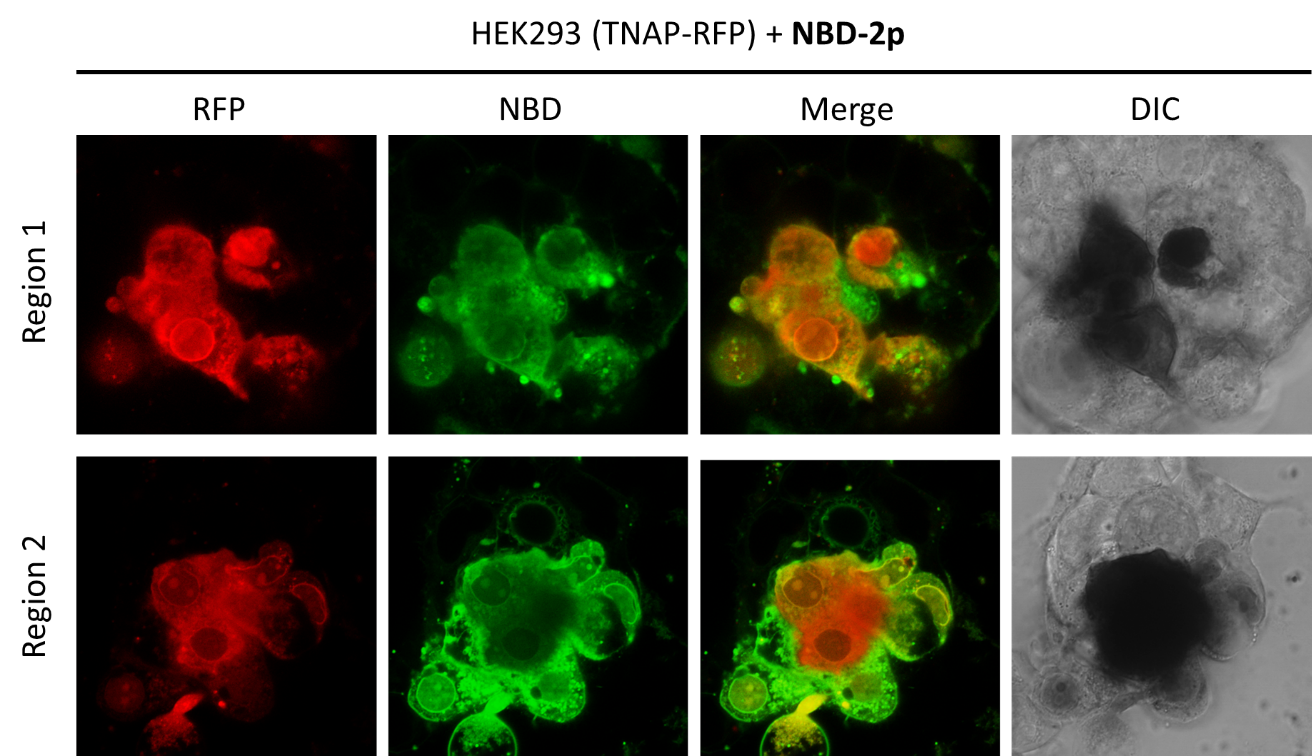

**Figure S6.** Confocal fluorescence imaging of HEK293 cells, after the transfection of TNAP-RFP gene, incubated with NBD-2p (200 μM) for 4 h.

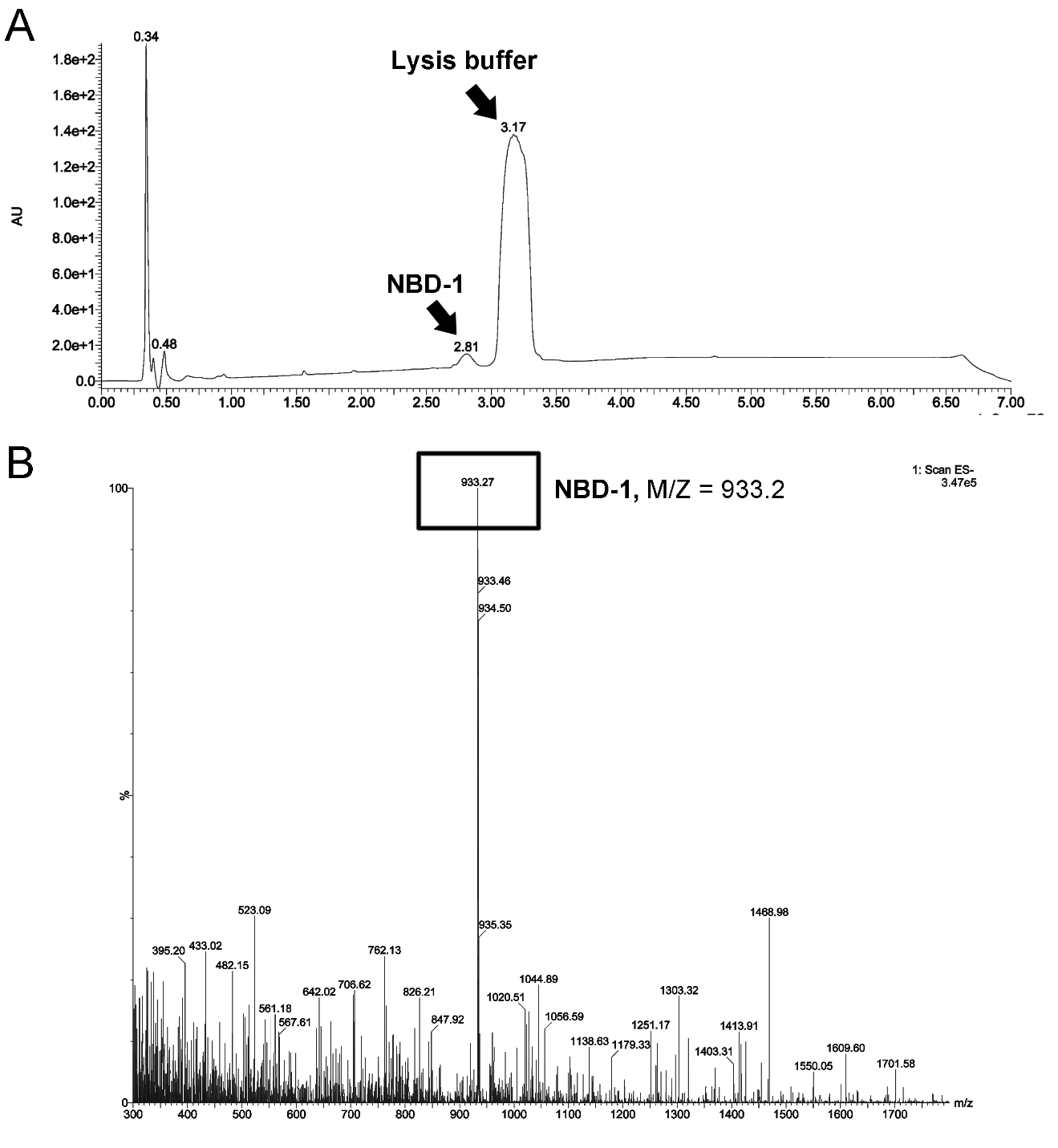

**Figure S7.** LC-MS analysis of cell lysate of HEK293 cells, after TNAP-RFP gene transfection, incubated with NBD-1p (200 μM, 24 h). (A) HPLC spectrum of cell lysate of HEK293_TNAP-RFP cells incubated with NBD-1p (200 μM, 24 h). (B) Mass spectrometry of the peak (retention time = 2.81 min) in (A). The dephosphorylation of the peptide is confirmed.

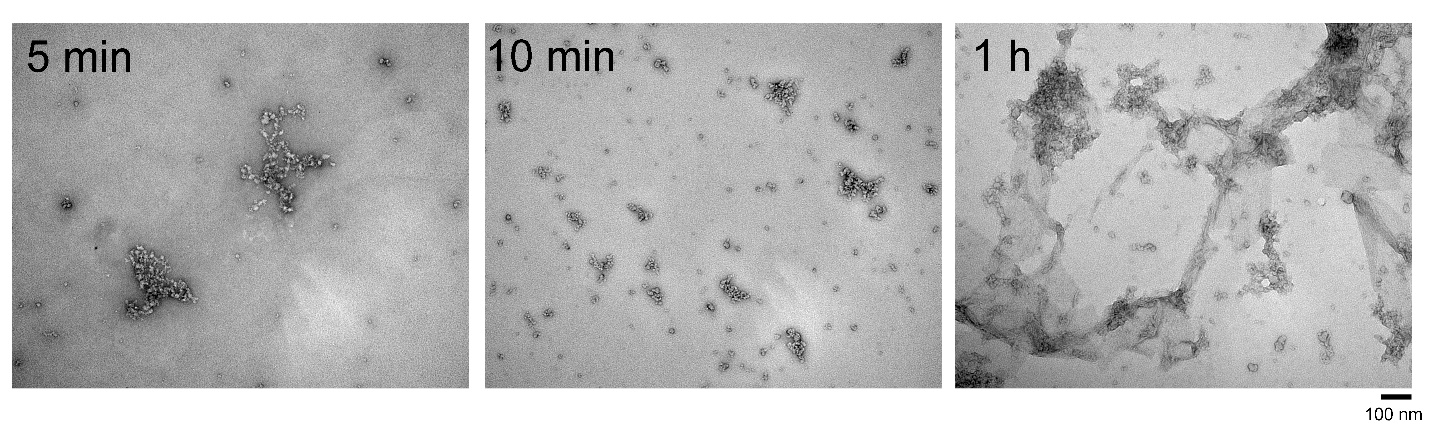

**Figure S8**. TEM images of NBD-1p (200 μM) incubated with ALP (0.1 U/mL) for different time.

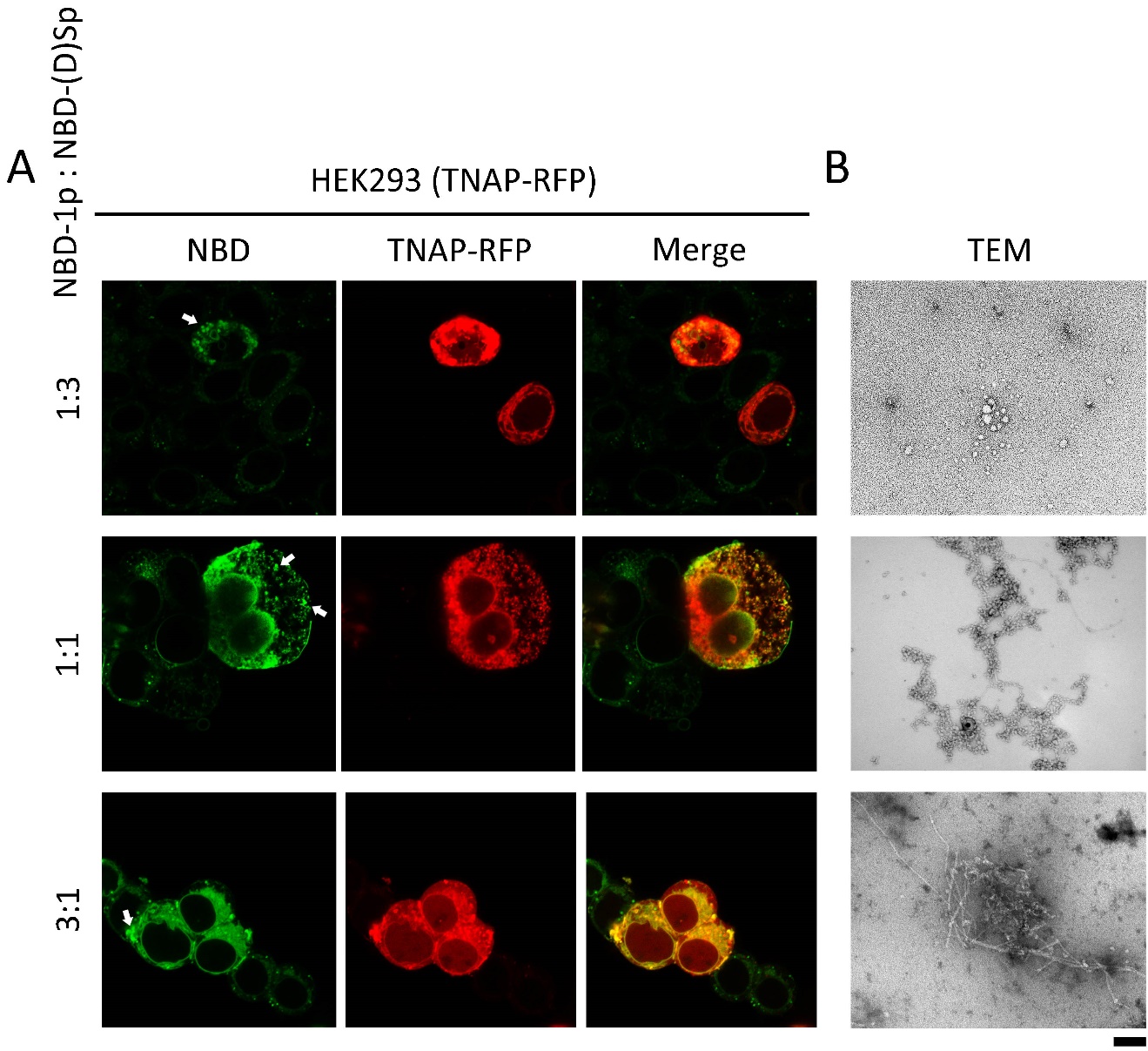

**Figure S9.** HEK293 cells, after TNAP-RFP gene transfection, incubated with the co-assemblies of NBD-1p and NBD-(D)Sp. (A) Confocal fluorescence images of HEK293_TNAP-RFP cells incubated with the mixture of NBD-1p and NBD-(D)Sp at the ratio of 1:3, 1:1, and 3:1, respectively. Endo/lysosomal retention is indicated by arrows. (B) TEM images of the mixture of NBD-1p and NBD-(D)Sp at the ratio of 1:3, 1:1, and 3:1, respectively, after the addition of ALP (0.1 U/mL, 3 h). Total peptide concentration is 200 μM.

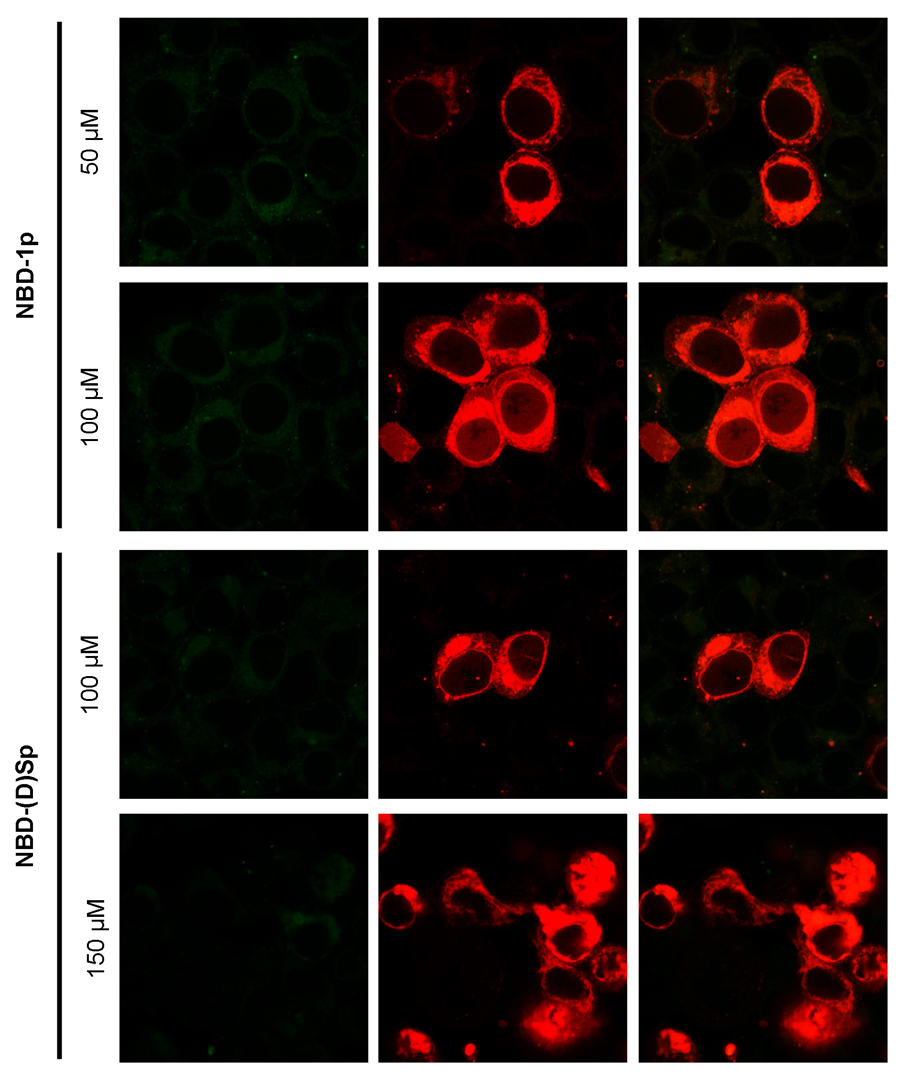

**Figure S10.** HEK293 cells, after TNAP-RFP gene transfection, incubated with NBD-1p (100 or 50 μM) or NBD-(D)Sp (100 or 150 μM) below CMC, respectively.

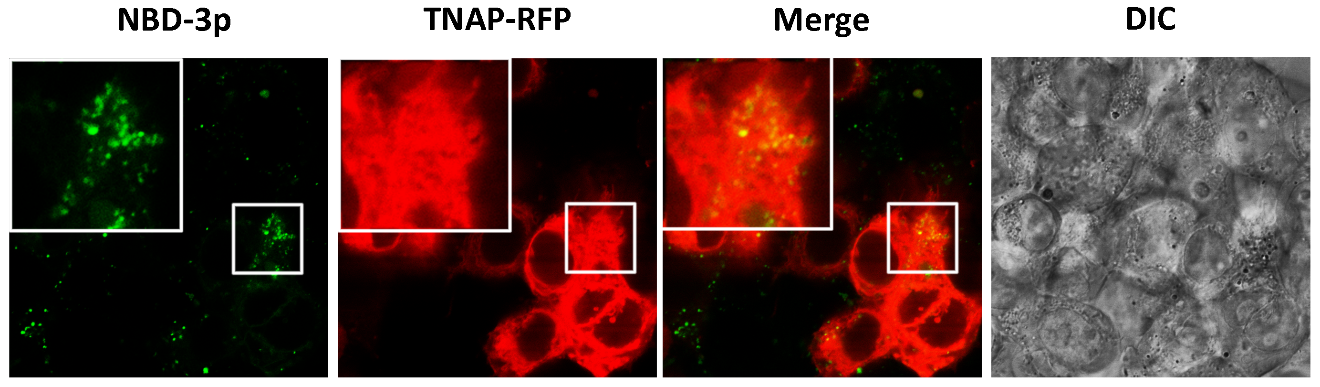

**Figure S11.** Confocal fluorescence imaging of HEK293 cells, after the transfection of TNAP-RFP gene, incubated with NBD-3p (400 μM) for 12 h. The fluorescent puncta indicate the endosomal retention of NBD-3p.

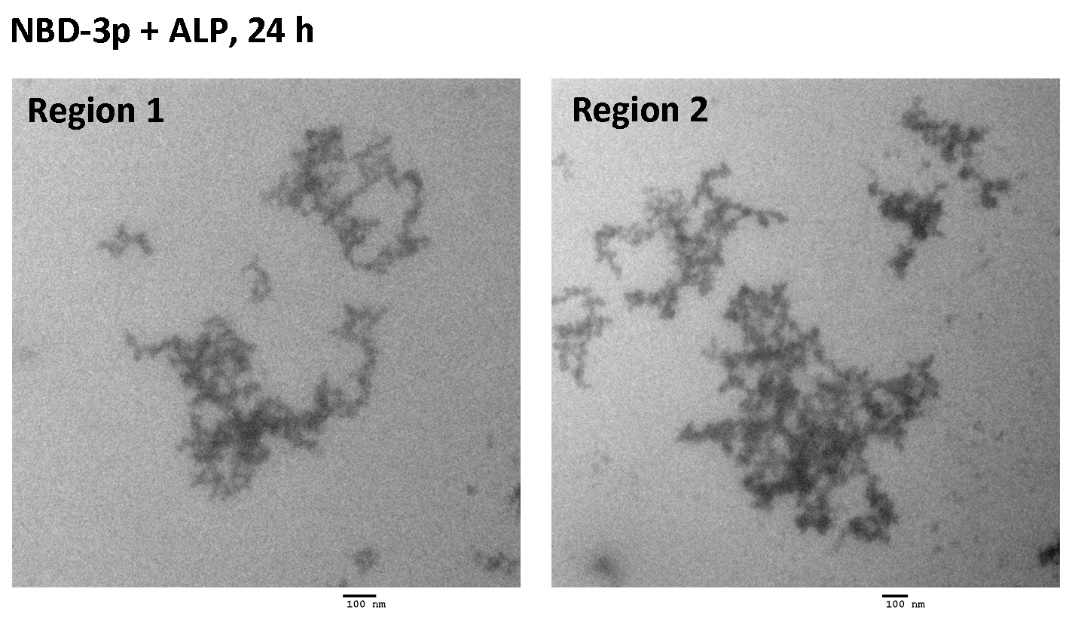

**Figure S12.** TEM images of NBD-3p (500 μM) incubated with ALP (1 U/mL, 37℃) for 24 h.

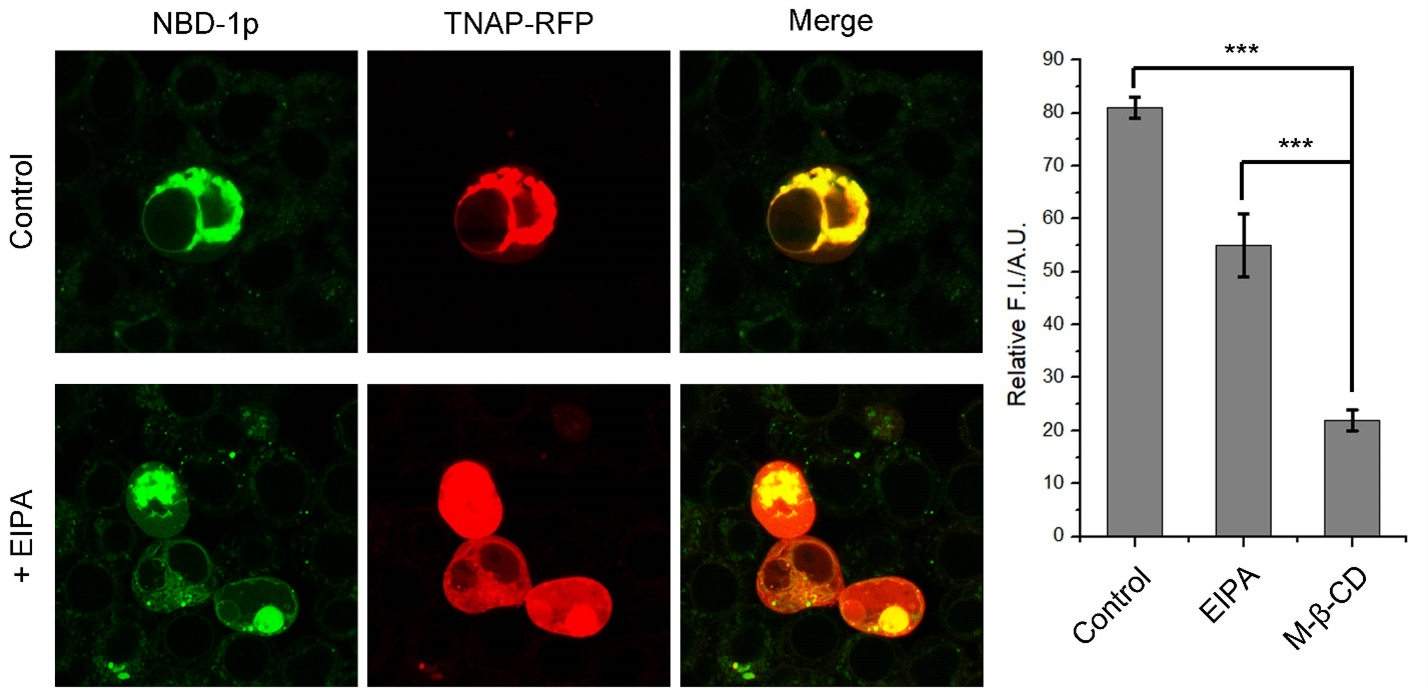

**Figure S13**. Confocal fluorescence images of HEK293_TNAP-RFP cells incubated with **NBD-1p** (200 μM, 12 h) in the presence of macropinocytosis inhibitor (5-[N-ethyl-N-isopropyl] amiloride, EIPA, 5 μM). *** means p<0.001.

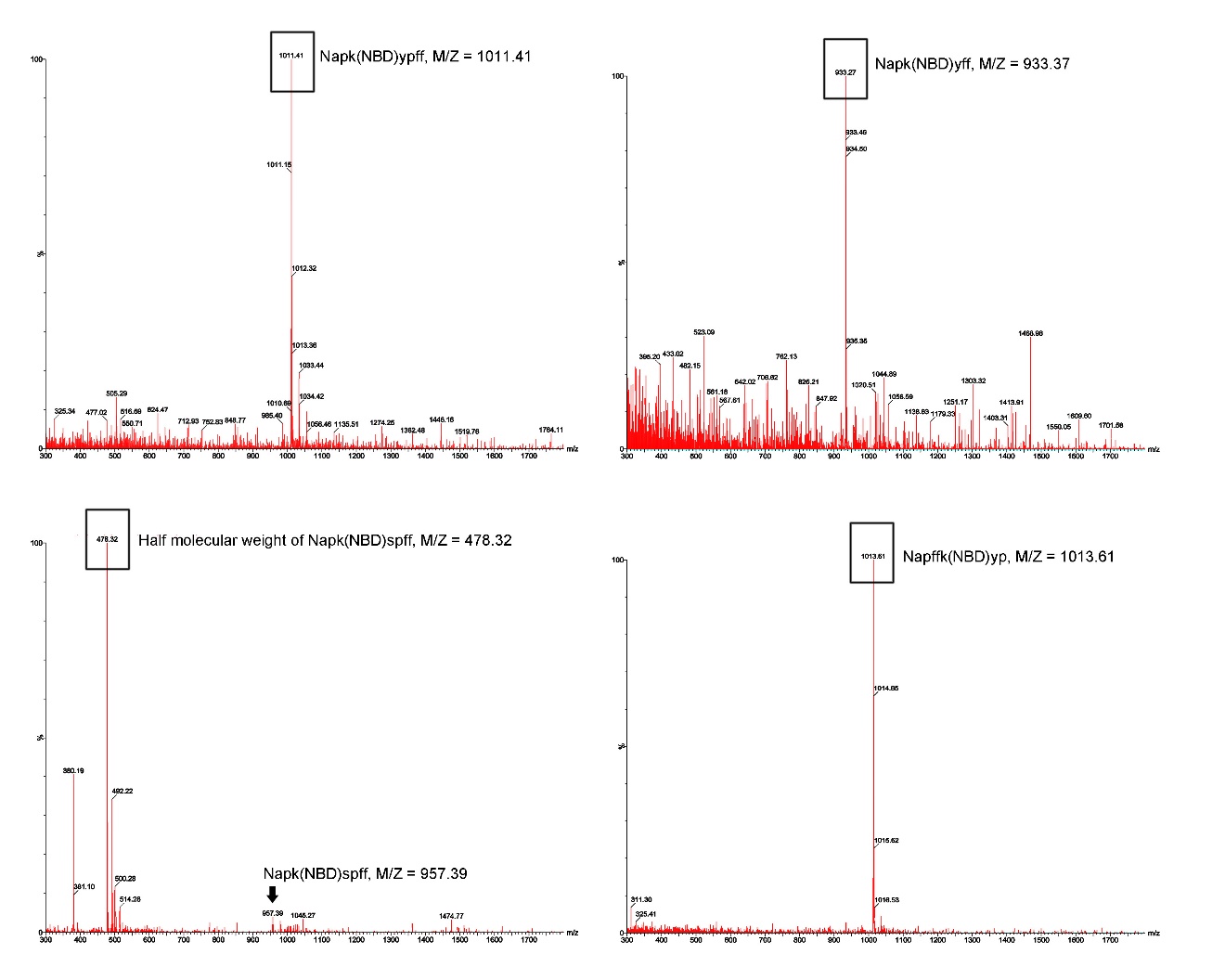

**Figure S14**. Mass spectrum of NBD-1p (Napk(NBD)ypff), NBD-(D)Sp (Napk(NBD)spff), NBD-1 (Napk(NBD)yff) and NBD-2p (Napffk(NBD)yp).
